# WISER: an innovative and efficient method for correcting population structure in omics-based selection and association studies

**DOI:** 10.1101/2025.02.07.637171

**Authors:** Laval Jacquin, Walter Guerra, Mariusz Lewandowski, Andrea Patocchi, Marijn Rymenants, Charles-Eric Durel, François Laurens, Maria José Aranzana, Hélène Muranty

## Abstract

This work introduces WISER (whitening and successive least squares estimation refinement), an innovative and efficient method designed to enhance phenotype estimation by addressing population structure. WISER outperforms traditional methods such as least squares (LS) means and best linear unbiased prediction (BLUP) in phenotype estimation, offering a more accurate approach for omics-based selection and association studies. Unlike existing approaches which correct for population structure, WISER offers a generalized framework that can be applied across diverse experimental setups, species, and omics datasets, such as single nucleotide polymorphisms (SNPs), near-infrared spectroscopy (NIRS), and metabolomics. Within its framework, WISER extends classical methods that use eigen-information as fixed-effect covariates to correct for population structure, by relaxing their assumptions and implementing a true whitening matrix instead of a pseudo-whitening matrix. This approach corrects fixed effects (e.g., environmental effects) for the genetic covariance structure embedded within the experimental design, thereby removing confounding factors between fixed and genetic effects. To support its practical application, a user-friendly R package named `wiser` has been developed. The WISER method has been employed in analyses for genomic prediction and heritability estimation across four species and 33 traits using multiple datasets, including rice, maize, apple, and Scots pine. Results indicate that genomic predictive abilities based on WISER-estimated phenotypes consistently outperform the LS-means and BLUP approaches for phenotype estimation, regardless of the predictive model applied. This underscores WISER’s potential to advance omics analyses and related research fields by capturing stronger genetic signals.

## 1. Introduction

In the realm of modern quantitative genetics, accurate phenotype estimation relies heavily on advanced statistical models that analyze the vast and complex data generated from experimental designs. These models are crucial for effectively capturing the nuances of phenotypic variation and ensuring reliable assessments of traits across diverse environments and conditions. A significant challenge emerges when population structure—defined as the presence of genetically different subgroups (i.e., unequal genetic covariances among individuals) within a sample—is not adequately addressed. Failure to consider population structure can confound the relationship between genetic variants and phenotypic traits (Byrne et al., 2020; Gloss et al., 2023; Sul et al., 2018). This oversight may ultimately lead to spurious associations and inaccurate estimates of genetic effects (Astle & Balding, 2009; Pritchard et al., 2000). Correcting for population structure is, therefore, a critical step in both genomic selection and genome-wide association studies (GWAS), as these approaches depend on accurate estimates of the genetic marker effects associated with traits of interest (Azevedo et al., 2017; Daetwyler et al., 2012; Kang et al., 2008; Patterson et al., 2006; Price et al., 2006; Riedelsheimer et al., 2012; Yu et al., 2006).

To address the confounding effects of population structure, numerous methodologies have been developed, in human, animal and plant genomics. Tools such as principal component analysis (Price et al., 2006) and linear mixed models (Kang et al., 2008) have been implemented to correct for unequal genetic relatedness within population in GWAS. Similarly, methods that incorporate eigen-information as fixed-effect covariates—derived from the spectral decomposition of kinship or genomic covariance matrices—within the framework of genomic best linear unbiased prediction (GBLUP) and its derivatives have proven essential in mitigating these confounding factors (Azevedo et al., 2017; Daetwyler et al., 2012; Riedelsheimer et al., 2012; Yang et al., 2011).

Effectively addressing population structure is particularly important in plant breeding populations, where variations in relatedness among individuals, arising from factors such as the sampling of genetic resources among others, can act as a significant confounding factor. For instance, in crops such as maize and rice, structured populations due to geographic origins or selective breeding practices can lead to biased estimates of marker-trait associations if population stratification is not adequately addressed (Zhao et al., 2011). Despite its potential impact, population structure is often overlooked, especially when estimated to be weak (Baertschi et al., 2021; Jung et al., 2020; Millet et al., 2019). In experimental designs, phenotypic estimation for each genotype is typically conducted using a least-squares (LS) means approach (Amadeu et al., 2020; Jung et al., 2020; Lorenzi et al., 2022, 2024; Njuguna et al., 2023; Rio et al., 2021; Sakhale et al., 2023; Samira et al., 2020; Tucker et al., 2022; Vogel et al., 2022), which does not account for the confounding effects of the genetic covariance structure among individuals in these designs. This oversight can introduce significant confounding factors, particularly related to fixed environmental effects, during the phenotype estimation process, ultimately resulting in less accurate estimates.

The best linear unbiased predictor (BLUP) is another commonly used method for phenotypic estimation per genotype, sometimes adjusting for population structure (Chen et al., 2022; Li et al., 2018; McLeod et al., 2023; Zhang et al., 2016). In this approach, principal components derived from PCA on omic data are often included as fixed-effect covariates to account for population structure. However, there is no consensus on the effectiveness of this method for consistently accounting for population structure, as findings have been mixed. Some studies argue that principal components may not reliably capture population structure (Carress et al., 2021; Elhaik, 2022; Lawson et al., 2020; Liu et al., 2013; Yao & Ochoa, 2023), noting limitations such as subjective selection of the number of components and low variance explained by the first few components in omic data, which may poorly represent the data structure (Carress et al., 2021; Elhaik, 2022). Additionally, in some association studies, principal component correction for population structure has been shown to fail in controlling the false positive rate (Elhaik, 2022; Liu et al., 2013) and can reduce the detection of true positives (Elhaik, 2022; M. Wang et al., 2021).

In response to this need, we propose WISER (whitening and successive least squares estimation refinement), a novel approach specifically designed to efficiently handle the genetic covariance structure between individuals within experimental designs. Unlike previous methods, which are often restricted to specific experimental setups or species (e.g., animals), WISER offers a more generalized framework that can be applied across diverse experimental designs (e.g., alpha lattice, multi-environment trial, multi-environment provenance-progeny trial, etc.), species, and omics datasets—including single nucleotide polymorphisms (SNPs), near-infrared spectroscopy (NIRS), and metabolomics. This versatility makes it particularly useful in agricultural research, where multiple experimental setups exist and different types of omics data are often combined to enhance the accuracy of omics-based selection and marker-trait associations.

Another feature that distinguishes WISER from existing methods is its ability to extend the use of eigen-information as fixed-effect covariates, which is the most common method for correcting population structure (Azevedo et al., 2017; Daetwyler et al., 2012; Riedelsheimer et al., 2012; Zhou & Stephens, 2012). By whitening fixed-effect variables, WISER effectively corrects for the genetic covariance structure among individuals in an experimental design, thereby minimizing confounding effects and enabling more accurate phenotype estimation. This is particularly important in plant breeding programs, where the ability to accurately conduct omics-based selection or associate markers with traits under selection is important for developing new crop varieties that exhibit improved yield, enhanced disease resistance, and greater environmental adaptability (Y. Xu & Crouch, 2008).

Additionally, WISER is implemented as a user-friendly R package, ensuring accessibility and practicality for researchers and practitioners in the field. The package includes intuitive tools that facilitate the automated optimization of key parameters, such as the selection of whitening matrices, which are crucial for effectively correcting population structure. This automation streamlines the estimation process, minimizing the risk of user error and enhancing the method’s applicability across diverse research environments. Furthermore, WISER’s versatility extends beyond plant applications; it has the potential to be used with longitudinal datasets in animal breeding programs, thereby broadening its utility across the agricultural sciences (Henderson, 1984; VanRaden, 2008).

In addition to its computational accessibility, WISER is designed to integrate seamlessly with existing pipelines for omics-based selection and GWAS, providing a comprehensive solution for researchers looking to improve the accuracy and reliability of their analyses. By offering a method that corrects for population structure across multiple experimental setups, species and omics data types, WISER enhances the ability to perform integrative analyses, where insights from genomics, transcriptomics, and metabolomics can be combined to paint a more complete picture of the genetic architecture underlying complex traits (Hu et al., 2021).

The rest of this paper is organized as follows: Section 2.1 presents the WISER framework, explaining how its whitening process aims to effectively remove the confounding effects of population structure on fixed-effect variables, thereby improving phenotype estimation.

Section 2.2 demonstrates that previous approaches, particularly the covariance analysis via eigenvectors (EVG) approach introduced by Yang et al. (2011) and Azevedo et al. (2017), can be viewed as a specific case of the WISER framework, where WISER extends EVG by relaxing its underlying assumptions and implementing a true whitening matrix instead of a pseudo-whitening matrix. The EVG approach adjusts fixed-effect variables for population structure and has shown superior genomic predictive ability compared to several other methods, as highlighted by Azevedo et al. (2017). This provides a solid foundation for illustrating how WISER extends and enhances this methodology, as well as other models that use principal components (PC) as fixed effects. Furthermore, this section shows that the ordinary least squares (OLS) estimate of fixed effects in the EVG approach is identical to that in the PC model, under the relaxed assumptions, further emphasizing WISER’s broader extension to models that incorporate eigen-information as fixed effects.

Section 2.3 details the classical LS-means and BLUP methods used as benchmarks for comparison with WISER in phenotype estimation. For a fair comparison with WISER, BLUP values were computed by including principal components derived from PCA on genomic data as fixed-effect covariates. This section also details the predictive models employed to evaluate each phenotypic estimation method across diverse species and traits. Additionally, it describes how genomic heritability estimates were calculated for phenotypes derived from the WISER, LS-means, and BLUP methods. Finally, it outlines different computed statistics aimed at explaining the improved genomic predictive ability when using phenotypes derived from WISER.

Section 3 (Results) provides a comparative analysis of the genomic predictive abilities and heritabilities associated to phenotypes derived from WISER, LS-means, and BLUP, along with an exploration of the factors driving the differences in these results. In this section, we demonstrate that WISER’s correction for the genetic covariance structure between individuals in an experimental design enhances genomic predictive abilities and heritability estimates, thereby improving genomic selection and association studies through the capture of stronger genetic signals.

The R package `wiser` can be easily installed from GitHub at https://github.com/ljacquin/wiser. To fully align with the FAIR principles, all repositories containing the datasets and R scripts used in this study’s analyses are publicly available on GitHub: https://github.com/ljacquin?tab=repositories.

## 2. Materials and methods

### 2.1 The WISER statistical framework

In the following, we will exclusively use the term “genotype”, though it can be interchangeably replaced with “animal” in other contexts such as longitudinal data analysis. Let *u*= (*u*_1_, …, *u*_*q*_)′represents the vector of genetic values associated with the *q* genotypes, where 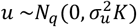 and *K* denotes the genetic covariance matrix between genotypes estimated from omic data. For an experimental design, the WISER statistical framework solves the following model to estimate a vector ν = (ν_1_, …, ν_*q*_)′of phenotypes which is a proxy for the vector *u*of genetic values:

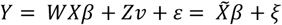

where:

- *Y*(*n* x 1) is the vector of individual phenotypic measurements, with values repeated for each genotype (or animal) across one or more environments.
- *W* (*n* x *n*) is a whitening matrix built using omic data (e.g., SNP markers, metabolites, or reflectance wavelengths) that whitens the fixed-effect variables to account for the genetic covariance structure among individuals.
- *X*(*n* x *l*) is the design matrix that links fixed effects to individual phenotypic measurements.
- *β*(*l* x 1) is the vector of fixed effects.
- ν (*q* x 1) is the vector of *q* phenotypes, approximating the genetic values associated with *q* genotypes or animals and treated as a vector of fixed effects.
- *Z*(*n* x *q*) is the design matrix that links the *q* phenotypes to the individual phenotypic values in the experimental design.
- *ε* (*n* x 1) is the vector of residuals, where 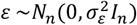.
- 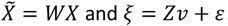.

Note that ν is modeled as a distinct vector of fixed effects with respect to *β*. Simultaneously, the whitening matrix *W* plays a pivotal role in this framework. It transforms the fixed-effect variables to remove the confounding effects of genetic covariance among individuals within the experimental design. In this context, 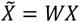 represents the whitened fixed-effect variables, enabling an effective correction for population structure. Let 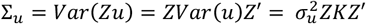 represents the genetic covariance matrix between individuals in the experimental design. For any covariance matrix Σ_*u*_, there exists a non-unique whitening matrix *W* that possesses the following property (Kessy et al., 2018; Podosinnikova, 2016; C. Wang et al., 2024):

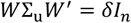

where *δ* is a strictly positive constant usually equal to one. There are several methods for computing *W*, including zero-phase component correlation analysis (ZCA-cor), principal component correlation analysis (PCA-cor) and cholesky (Cholesky) among others. Define 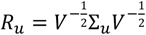 as the genetic correlation matrix associated with Σ_*u*_, where *V* = *diag*(Σ_*u*_) is a diagonal matrix with the variances of Σ_*u*_on its diagonal. Since Σ_*u*_and *R*_*u*_are symmetric and positive semi-definite, they have the following spectral decompositions: Σ_*u*_= *U*Λ*U*′and *R*_*u*_= *G*Θ*G*′, where *U* and *G* are orthogonal eigenvector matrices for Σ_*u*_and *R*_*u*_, respectively, while Λ and Θ are diagonal matrices of positive eigenvalues for Σ_*u*_and *R*_*u*_, respectively. The inverse square root matrices of Σ_*u*_and *R*_*u*_are given by 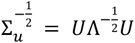 and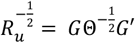, these satisfy 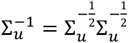 and 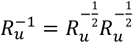 since *U* and *G* are orthogonal matrices. According to Kessy et al. (2018), the whitening matrices associated to ZCA-cor, PCA-cor and Cholesky are given by:

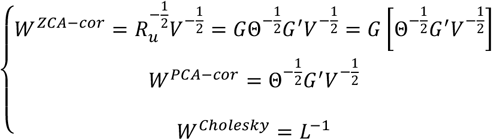

where *L* is a full-rank lower triangular matrix derived from the following Cholesky decomposition Σ_*u*_= *Var*(*Zu*) = *LL*′. The PCA-cor whitening procedure can be seen as standardizing variables using 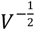, followed by a rotation using the transposed correlation eigenmatrix *G*′, and then scaling using the inverted correlation singular values matrix 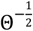. ZCA-cor whitening extends this by applying an additional rotation *G* to revert to the original basis of the standardized variables. While Kessy et al. (2018) recommend both ZCA-cor and PCA-cor, each whitening method is optimal under distinct criteria. WISER implements these two approaches alongside the Cholesky procedure. Notably, ZCA-cor uniquely ensures that the whitened variables maintain the highest correlation with the original variables. Kessy et al. (2018) provide a detailed discussion of these criteria and the optimality of each method.

These whitening matrices effectively eliminate any genetic covariance structure among individuals in an experimental design when such structure is present. A simple and clear example is provided by setting *W* = *L*^−1^and defining the vector *T* = (*t*_1_, …, *t*_*n*_)^′^= *Zu*, where *T* represents the individual genetic values in the experimental design. In this case, the left multiplication of *T* by *W* eliminates the genetic covariance structure between individuals from *T*, as illustrated by its covariance matrix:

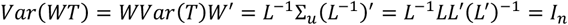

Although fixed effects are not inherently random, the whitening approach remains effective when significant confounding relationships exist between these effects and individual genetic values. This can be pedagogically illustrated through the following uncommon yet instructive case study, where a vector *F* of individual fixed effects, associated with a fixed-effect variable or factor, is highly correlated with *T*.

For instance, in an experimental design with *J* environments, where the environment variable is generally treated as a fixed-effect factor, we can express 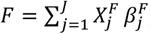, where 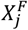 is the incidence column (composed of 0s and 1s) for the *j*^*th*^ environment, and 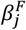 is the effect associated with this environment. If each genotype is uniquely associated with a specific environment, and genotype-environment interactions are considered negligible, a high correlation may arise between *T* and *F*, resulting in *F* ≈ *kT*, where *k* is a constant. In such scenarios, the variances and covariances of environmental effects may closely—or at least partially—mirror those of genetic effects, creating confounding associations. As a result, applying *W* to *F* effectively whitens the individual environmental effects with respect to the mirrored genetic covariance structure embedded in the experimental design. This process removes confounding effects between environments and genotypes, thereby preserving the integrity of the fixed-effect estimates for the environments.

Note that constructing the whitening matrix *W* depends on 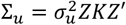, which, in turn, requires estimating the variance component 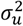. In WISER, this estimation, along with that of 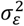, is performed using a parallelized approximate Bayesian computation (ABC) algorithm to ensure fast and stable variance component estimation for large datasets associated with complex experimental designs. Additionally, because Σ_*u*_may approach singularity in practical applications, WISER incorporates established shrinkage and regularization procedures to maintain positive-definiteness (Kwan, 2011; Nikovski & Byadarhaly, 2016; Schäfer & Strimmer, 2005; Steland, 2018; Theiler, 2012; Touloumis, 2015). Further details on the ABC algorithm and regularization procedures are provided in the Supplementary file of the Supplementary materials.

To automate the selection of whitening procedures and regularization parameters, WISER also includes a dedicated optimization function. This function performs a grid search across predefined combinations of whitening methods, regularization parameters, and various prediction methods, such as genomic best linear unbiased prediction (GBLUP), least absolute shrinkage and selection operator (LASSO), support vector regression (SVR), reproducing kernel Hilbert space (RKHS) regression, and random forest. Using k-fold cross-validation (CV), the function evaluates the performance of each combination and identifies the optimal whitening method and regularization parameter that maximize predictive ability (i.e., Pearson’s correlation), based on a majority vote across the k-fold CV results for a subset of the dataset.

In WISER, estimates for *β* and ν are obtained using the following successive OLS procedure:

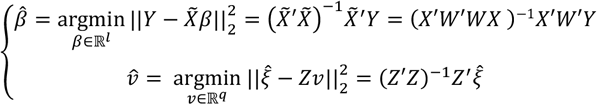

where 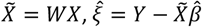 and 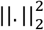 represents the squared *l*^2^ norm. Note that 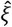 corresponds to the part of *Y* that is not explained by the transformed fixed-effect variables, which incorporate information derived from the omic data, and is orthogonal to these variables. As with any linear model, this orthogonality arises because 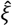 is estimated by projecting *Y* onto the space orthogonal to that spanned by the whitened fixed-effect variables, ensuring orthogonality by design:

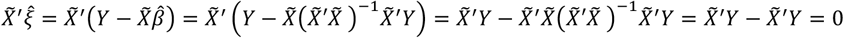

Thus, the vector 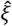 can be used to estimate the vector ν of phenotypes, serving as an approximation of the genetic values of the genotypes in omic-based selection and association studies. It is also important to note that 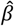 is computed using *W*, which is derived from Σ_*u*_and thus depends on an estimate of 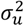. Furthermore, when no repeated measurements per genotype are available, the genetic covariance matrix Σ_*u*_associated with individuals in the experimental design becomes equivalent to the genetic covariance matrix *K*. In this case, the population structure is effectively represented by *K*. To mitigate issues of numerical singularity in certain practical applications, WISER also incorporates the Moore-Penrose generalized pseudo-inverse method for estimating *β* and ν.

### 2.2. WISER as an extension of models using eigen-information as fixed-effect covariates to correct for population structure

We recall that the spectral decomposition of Σ_*u*_can be expressed as follows (see Section 2.1):

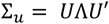

where Λ is the diagonal matrix of positive eigenvalues, and *U* is the orthogonal matrix of eigenvectors associated to the genetic covariance matrix. According to Azevedo et al. (2017), the EVG model which accounts for population structure correction is defined as follows:

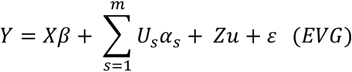

where (*U*_*s*_)_1≤*s*≤*m*_ represents the first *m* ≤ *n* eigenvectors, with the highest eigenvalues, of the orthogonal matrix *U* of eigenvectors associated to the genetic covariance matrix Σ_*u*_, and (*α*_*s*_)_1≤*s*≤*m*_ their corresponding effects. To align the notations of this model with those of the WISER approach, particularly with respect to the spectral decomposition of Σ_*u*_, we assume *m* = *n*. In this case, the EVG model can be rewritten as:

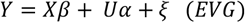

where the matrix *U* = (*U*_1_ … *U*_*n*_) is the column concatenation of the *n* eigenvectors (i.e., eigenvector matrix of Σ_*u*_), *α* = [*α*_1_, …, *α*_*n*_]^′^and ξ = *Zu*+ *ε*. Note that WISER also implicitly assumes that ξ = *Zu*+ *ε*, but in its framework, *u*is approximated by ν, for which no distributional assumption is made.

Let *E*_*U*_ = *span*(*U*) represent the vector subspace generated by the column vectors of *U*. Let *P*_*E*_ = *U*(*U*^′^*U*)^−1^*U*′ and 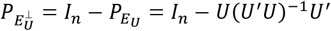 be the orthogonal projectors onto 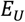 and 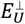 respectively, where 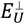 is the orthogonal complement of *E*_U_ (i.e.,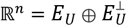). Using the ollowing linear transformation based on 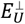, we have:

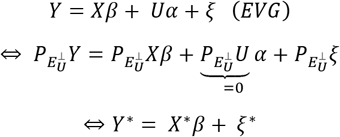

where 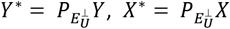 and 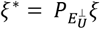. Note that (*U*_*s*_)_1≤*s*≤*n*_ ∈ *E*_*U*_, therefore we have 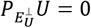. The last property can also be verified directly as follows:

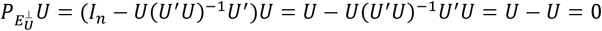

In the absence of assumptions regarding ξ^∗^, as is the case within the WISER framework, the OLS estimate for *β* in the *EVG* model is given by:

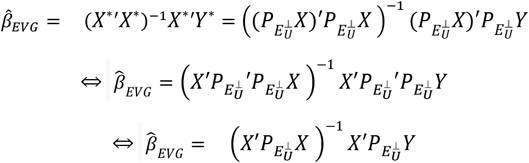

Note that 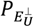 is symmetric and idempotent, therefore we have 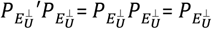. From the expression for 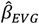, one can notice that the vectors of the design matrix *X* for fixed effects are orthogonally projected onto 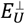 (i.e.,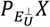), which represents the orthogonal complement of *E*_*U*_. This projection serves to eliminate the genetic covariance structure captured by the eigenvectors *U*_*s*_ (1 ≤ *s* ≤ *n*) from *X*, under the assumption that all principal axes of variation are considered, which, however, is typically not the case in practice, where only a fraction of the genetic covariance is accounted for. Furthermore, within the WISER framework, the projector 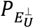 can be interpreted as a pseudo-whitening matrix 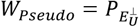 associated with its OLS estimate of *β*. Indeed, we recall that the OLS estimate of *β* related to WISER is expressed as follows:

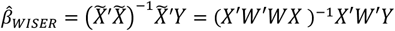

By substituting *W* with *W*_*Pseudo*_, we see that:

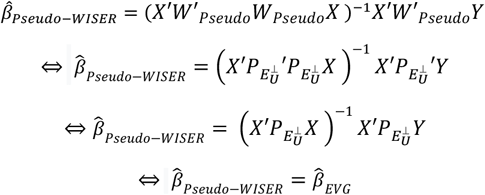

Therefore, 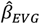 can be viewed as a specific case of the OLS estimate of *β* within the WISER framework, where a pseudo-whitening matrix *W*_*Pseudo*_ is used in the absence of assumptions regarding ξ^∗^.

We can show that *W*_*Pseudo*_ does not verify the whitening property. Indeed, for the vector *T* = (*t*_1_, …, *t*_*n*_)^′^= *Zu*∼*N*_*n*_(0, Σ_*u*_) we have:

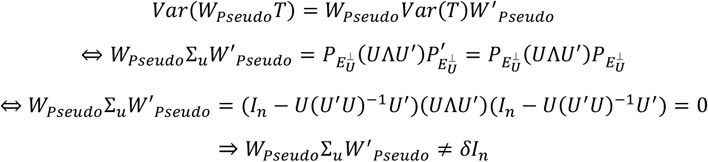

Thus, applying *W*_*Pseudo*_to whiten *T* results in a null covariance matrix, rather than an isotropic covariance matrix (i.e., proportional to the identity matrix), with zero variances along the diagonal. Consequently, using *W*_*Pseudo*_ to account for population structure does not properly whiten the fixed-effect variables in the experimental design. Nevertheless, *W*_*Pseudo*_ can be considered a pseudo-whitening matrix, as it serves as a rough approximation to a true whitening matrix. In fact, a true whitening matrix *W*_*Adj*._ can be derived from 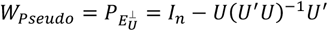 by adjusting the term *U*(*U*^′^*U*)^−1^*U*^′^as follows:

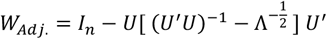

where the diagonal matrix 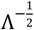 corresponds to the inverse of the singular values (i.e., square roots of the eigenvalues) of Σ_*u*_. We can verify the whitening property for *W*_*Adj*._ as follows:

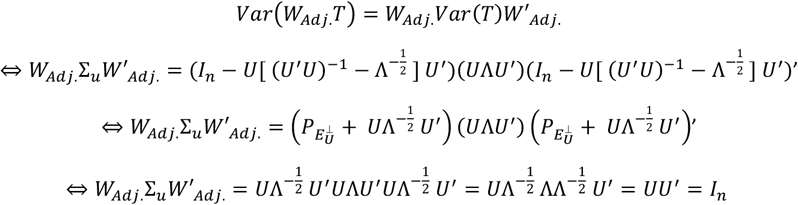

Therefore, *W*_*Pseudo*_ can indeed be regarded as a pseudo-whitening matrix, as it requires an adjustment with 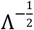 to be converted into the true whitening matrix *W*_*Adj*._.

The equality between the OLS estimates of the fixed effects in the EVG and PC models can be demonstrated as follows. According to Azevedo et al. (2017), the PC model, which corrects for population structure, is defined as follows:

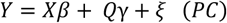

In this model, *γ* represents the regression coefficients associated with the PCs, and each column of the matrix *Q* corresponds to the PC coordinates of individuals along an axis directed by an eigenvector of *U*. We assume that all PCs are included in *Q* to ensure that the OLS estimates of the fixed effects are identical for both the PC and EVG models. In Azevedo et al. (2017), the matrix of PC coordinates is computed as *Q* = *UG*, where *G* corresponds to Σ_*u*_as the estimated genetic covariance between individuals, and in terms of dimensionality (i.e., both *G* and Σ_*u*_have the same number of rows and columns, equal to the number of individuals). This computational approach deviates from the standard method for computing PC coordinates. Nevertheless, in the context where PC coordinates are generated by projecting genotype data onto the axes defined by the eigenvectors of *U*, each column of *Q* must reside within the subspace *E*_*U*_. Consequently, we have 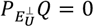. Therefore, applying a linear transformation based on 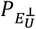 to the PC model, as implemented in the EVG model, results in 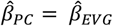.

### 2.3 Analyzed datasets, compared phenotypic estimation methods, predictive models, and computed statistics

#### 2.3.1. Analyzed datasets

Four datasets from experimental designs involving rice (Baertschi et al., 2021), maize (Millet et al., 2019), apple (Jung et al., 2022), and Scots pine (Perry et al., 2022) were used to compare genomic predictive ability (PA) and heritability estimation across 33 traits. The rice dataset contained 4,308 individual phenotypic measurements and 9,929 SNPs associated with 334 genotypes. The maize dataset included 18,970 individual phenotypic measurements and 41,723 SNPs associated with 246 genotypes. The apple reference population (REFPOP) dataset, after preprocessing to remove phenotypic outliers, consisted of 42,522 individual phenotypic measurements and 303,240 SNPs associated with 534 genotypes. Among these SNPs, only 50,000 were used, selected through uniform sampling to reduce computation time and resource usage, as Jung et al. (2020) demonstrated that a PA plateau is typically reached with as few as 10,000 SNPs for this dataset. Additionally, the apple dataset included two traits—scab and powdery mildew—not covered in Jung et al., 2022. Finally, the Scots pine dataset had 1,248 individual phenotypic measurements and 20,796 SNPs associated with 208 genotypes.

The traits predicted and analyzed for each species were as follows:

- Rice: days to flowering (FL), plant height (PH), grain yield (YLD), and grain zinc concentration (ZN).
- Maize: anthesis, anthesis-silking interval, ear height, grain number, grain weight, grain yield, plant height, silking, and tassel height.
- Apple: color over, flowering start, flowering end, full flowering, flowering intensity, fruit number, single fruit weight, fruit weight, harvest date, powdery mildew, russet frequency, scab, trunk diameter, and trunk increment.
- Scots pine: height (H), annual height increment (I), duration of budburst (D)—the time taken to progress from stage 4 (scales open along the length of the shoot, no needles) to stage 6 (green-tipped needles visible), and time taken to reach individual budburst stages: stage 4 (T4), stage 5 (T5), and stage 6 (T6).

For each dataset, the environmental variable was defined as a combination of factors, including site (or country), year, management type, block, and genotype generation, depending on the variables available. In the rice dataset, the environment was characterized by the combination of site, year, block, and genotype generation. For maize, it was defined by site, year, management type (rainfed or irrigated), and block. For apple, the environmental variable was defined as the combination of country, year, and management type (standard irrigation and pesticide use, standard irrigation with reduced pesticide use, or reduced irrigation with standard pesticide use). Finally, in the case of pine, the environment was determined by site, year, and block.

#### 2.3.2. Compared phenotypic estimation methods

To account for spatial variation prior to phenotypic estimation using the LS-means and BLUP approaches, a spatial heterogeneity correction was applied to each combination of trait and environment, resulting in spatially adjusted individual phenotypes. This correction was implemented when spatial location information was available, which was the case for the maize, apple, and pine datasets, where row and column, row and position, and GPS coordinates were respectively available. The spatial heterogeneity correction was performed using the spatial analysis of field trials with splines method (Rodríguez-Álvarez et al., 2018), following the same approach as described in Jung et al. (2020).

For the LS-means approach, the following multiple linear regression model was initially fitted:

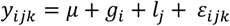

where *y*_*ijk*_ was the *k*^*th*^ spatially adjusted individual phenotype associated to genotype *i* in environment *j, g*_*i*_ was the fixed effect of genotype *i* (1 ≤ *i* ≤ *q*), *μ* was the fixed overall mean, *l*_*j*_ was the fixed effect of the *j*^*th*^ environment, and *ε*_*ijk*_ was the error term. The error term *ε*_*ijk*_ was assumed to follow a normal distribution, 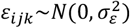, with its associated vector *ε* considered to follow a multivariate normal distribution with an isotropic covariance matrix. Phenotypic LS-means for the genotypes were then computed across all environments using the R package `lsmeans` (Lenth, 2016) to estimate a single phenotype value per genotype.

For the BLUP approach, the following linear mixed model, which accounts for population structure, was fitted using the R package `lme4` (Bates et al., 2015):

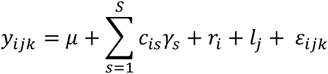

In this model, *y*_*ijk*_, *μ, l*_*j*_ and *ε*_*ijk*_ were defined as in the LS-means approach. Here, *r*_*i*_ represents the random effect of genotype *i* (1 ≤ *i* ≤ *q*), with 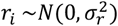, and its associated vector *r* was assumed to follow a multivariate normal distribution with an isotropic covariance matrix. Additionally, principal component (PC) coordinates for each genotype *i*, denoted *c*_*is*_ (1 ≤ *s* ≤ *S*) and treated as fixed effects, were computed using genomic data from each dataset with the R package `mixOmics` (Rohart et al., 2017). In this context, *γ*_*s*_ represented the effect associated with the *s*^*th*^ PC. Population structure is generally regarded as a low-dimensional process embedded within a high-dimensional space (Hoffman, 2013; Patterson et al., 2006). Consequently, a relatively small number of principal components, rarely exceeding 10, is assumed to effectively capture the underlying genetic structure of the population (Price et al., 2006). The number *S* of principal components (PCs) selected for computing the BLUP values of *r* was chosen to minimize the Akaike information criterion (AIC) within the range of values tested (1 to 10), as AIC is widely regarded as one of the most commonly used criteria for model comparison (Beugin et al., 2018).

Note that for the rice dataset, *y*_*ijk*_ corresponded to individual phenotypic measurements in both the LS-means and BLUP approaches, as spatial location data was unavailable. Additionally, in the LS-means approach, the Moore-Penrose generalized pseudo-inverse was used during the multiple linear regression procedure to address issues related to aliased variables (i.e., perfect multicollinearity) for certain traits. This adjustment was particularly necessary for traits D, H, and I in the pine dataset.

In the wiser framework, the overall mean (included by default) and the environmental variable were modeled as fixed-effect factors for the rice dataset. For the apple dataset, the overall mean, the environment (comprising site, year, and management), and rows and positions for spatial adjustment were modeled as fixed-effect factors, with rows and positions treated as environment-specific factors. For the maize and pine datasets, the overall mean was included as a fixed-effect factor, while row and column numbers (for maize) and GPS coordinates (for pine) were treated as fixed-effect quantitative variables specific to each environment. This approach minimized the number of fixed-effect variables, thereby reducing the risk of inestimability. For example, the maize dataset had 1,061 levels for the environment variable related to anthesis, resulting in 2,122 fixed-effect quantitative variables when accounting for row and column within each environment. Thus, the number of fixed-effect variables would have been considerably larger if row and column numbers had been modeled as factors within each environment. The three whitening procedures implemented in `wiser` are based on the R package `whitening` (Korbinian Strimmer, Takoua Jendoubi, Agnan Kessy, Alex Lewin, 2018). For each trait, the whitening procedure and regularization parameter were automatically selected using the `optimize_whitening_and_regularization` function from the `wiser` package, which was specifically designed for automated selection. As described in Section 2.1, this function performs a grid search over a subset of the dataset to identify the optimal combination of whitening procedure and regularization parameter that maximizes predictive ability (PA).

#### 2.3.3 Compared predictive models

Five predictive models were employed for genomic prediction: random forest (RF), support vector regression (SVR), genomic best linear unbiased prediction (GBLUP), reproducing kernel Hilbert space (RKHS) regression, and least absolute shrinkage and selection operator (LASSO). These models were selected to represent diverse genetic architectures and to evaluate whether the genomic predicted ability (PA) rankings across the different phenotypic estimation methods and traits remained consistent, regardless of the prediction approach. The RF, SVR, and LASSO models were fitted using the R packages `ranger` (Wright & Ziegler, 2017), `kernlab` (Karatzoglou et al., 2004), and `glmnet` (Friedman et al., 2010), respectively. GBLUP and RKHS regression were fitted using the R package `KRMM` (Jacquin et al., 2016). For RF, the number of randomly selected candidate variables (`mtry`) for testing splits at each tree node was set to one-third of the total number of SNPs, a common choice for regression tasks, and the number of trees was fixed at 1,000. For SVR, LASSO, GBLUP, and RKHS regression, hyperparameters were estimated or selected following the approach described in Jacquin et al. (2016). Specifically, for SVR, a Gaussian kernel was applied using the `ksvm` function, and its bandwidth parameter was estimated automatically using the heuristic in the `sigest` function. The regularization parameter *C* for SVR was computed as 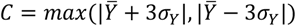, where 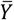 is the mean of phenotypes, as recommended by Cherkassky & Ma (2004). The *ε*-insensitive loss parameter in SVR was set to its default value of 0.1, as tested values ranging from 0.01 to 0.5 produced similar results, which were comparable to those of other prediction models. For LASSO, the regularization parameter was determined using K-fold cross-validation implemented in the `cv.glmnet` function, with the default value of `nfolds` set to 10. For GBLUP, the `krmm` function was applied with a linear kernel, using the default initial guess values for the variance component estimates required by the EM-REML algorithm. The same settings were used for RKHS regression, except that a Gaussian kernel was applied. The Gaussian kernel’s bandwidth parameter referred to as the rate of decay in `krmm` was set to its default value of 0.1, which provided comparable results to the other prediction models.

A 5-fold cross-validation (CV) scheme with 20 random shufflings of the datasets was employed to estimate the distributions of predictive ability (PA) and genomic heritability (*h*^2^) across various prediction models and phenotype estimation methods for all traits and species. For each shuffle, a new seed was generated by multiplying the shuffle number by 100, thereby introducing substantial variation in the randomized genotype distribution across shuffles. This approach generated a distribution of 100 estimated PA values for each combination of species, trait, prediction model, and phenotype estimation method. Similarly, a distribution of 100 estimated *h*^2^ values was generated with the GBLUP model using this approach. For each *k*^*th*^ validation fold, PA was assessed by calculating the Pearson correlation between the predicted phenotypes and those derived from the LS-means, BLUP, and WISER approaches. In contrast, *h*^2^ was computed from the estimated variance components of the following GBLUP model, which was fitted on the remaining *K* − 1 folds:

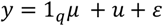

In this model, *y* = (*y*_1_, …, *y*_*q*_)′represents the vector of *q* phenotypes derived from either the LS-means, BLUP, or WISER phenotype estimation method. The vector 1_*q*_ = (1, …,1)^′^consists of *q* ones, and *μ* corresponds to the fixed overall mean. The vector *u*= (*u*_1_, …, *u*_*q*_)′represents the genetic values associated with the *q* genotypes, where 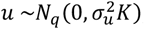, with *K* = *MM*^′^being the scaled genomic covariance matrix (i.e., the Gram matrix), and 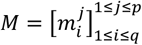 is the centered SNP marker matrix of size *q* x *p* with 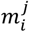 coded as 0, 1 or 2. The error vector *ε* = (*ε*_1_, …, *ε*_*q*_)′follows a normal distribution,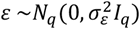. From this model, the genomic heritability *h*^2^ was defined as:

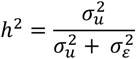

#### 2.3.4 Computed statistics

For the generated distributions of predictive ability (PA) and genomic heritability (*h*^2^), median values and deviations, computed across specific phenotype estimation methods, were calculated to facilitate graphical representations and subsequent analyses. To explore the relationships between PA and *h*^2^, deviations in the PA medians were compared to those in the *h*^2^ medians. Additionally, the deviations in the PA medians were compared to the Shannon entropy of genotype frequencies across all trait-species combinations. For each trait-species combination, the Shannon entropy of genotype frequencies was defined as:

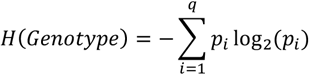

where *p*_*i*_ is the frequency of the *i*^*th*^ genotype, and *q* is the total number of genotypes. This comparison was intended to assess the impact of imbalanced genotype frequencies across all trait-species combinations on the deviations observed in the PA medians. A similar comparison was made between these deviations and the Shannon joint entropies of genotype and site, as well as genotype and environment. For each trait-species combination, these joint entropies were computed as follows:

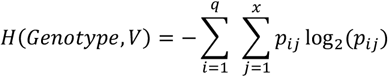

where *V* is either the site or environment variable, *p*_*ij*_ represents the joint frequency of genotype *i* and site (or environment) *j*, and *x* is the number of sites (or environments). These joint entropies were used to quantify the confounding degree between genotypes and their respective sites or environments.

To assess the degree of association between genotype and site or environment, and identify potential multicollinearity among their dummy variables, Cramér’s V statistic was computed. It was particularly reported when PA median values were low for a trait-species combination, helping to identify possible causes for these low values. This statistic was computed using the R package `vcd` (Meyer et al., 2002).

To further explore the deviations in PA medians, dimensionality reduction and clustering techniques were employed to evaluate genetic differentiation between genotypes. Uniform manifold approximation and projection (UMAP) (McInnes et al., 2018) was used to generate five-dimensional representations of the genomic data across all species. The choice of five dimensions was a trade-off between preserving enough genetic variability for effective clustering and mitigating the “curse of dimensionality”, which can reduce the accuracy of distance-based methods in higher-dimensional spaces. The UMAP approach was implemented using the R package `umap` (Konopka, 2023). The resulting five-dimensional representations were clustered using the *K*-means algorithm for each trait-species combination, with the optimal number of clusters determined as the value of *K* that maximized the Calinski-Harabasz (CH) index. The CH index was evaluated over a range of *K* values from 2 to 30 and is defined as:

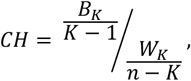

where:

- 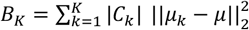 is the between-cluster sum of squares (BCSS) for the SNP marker data. Here, 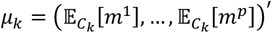 represents the centroid vector for cluster *C*_*k*_, *μ* = (𝔼[*m*^1^], …, 𝔼[*m*^*p*^])^′^is the mean vector for the entire dataset, and |*C*_*k*_| is the number of data points (i.e., vectors) in cluster *C*_*k*_.
- 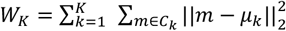 is the within-cluster sum of squares (WCSS), quantifying the compactness of data points around their respective cluster centroids. Here, *m* = (*m*^1^, …, *m*^*p*^)′represents an SNP vector with values coded as 0, 1 or 2.

The CH index generalizes Fisher’s test statistic from a one-way ANOVA with *K* levels to the multidimensional setting. While this index is not a test statistic and lacks a formal statistical distribution, it serves as a robust metric to balance the compactness of clusters with their separation, providing an effective measure for determining the optimal clustering structure. Note that *K*-means clustering aims to find the optimal partitioning of data into *K* clusters that minimizes *W*_*K*_. Therefore, the CH index is a natural choice for selecting *K*, as both the CH index and *K*-means are related to the total sum of squares (TSS) decomposition, which is expressed as:

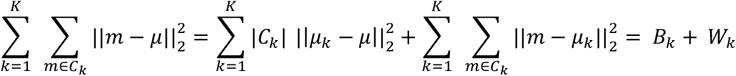

The CH index was computed using the R package `fpc` (Hennig, 2003). Finally, to quantify genetic differentiation between genotypes, the average fixation index *F*_*ST*_ was computed using genomic data associated with the *K*-estimated clusters for each trait-species combination. Notably, for each trait-species combination, the genomic data used accounted for genotype incidence relative to the available phenotypic measures in the experimental design. To improve computational efficiency and minimize resource usage, a stratified sampling approach was employed to maintain the data structure, while limiting the genomic dataset to 5,000 rows per trait-species combination. The *F*_*ST*_ values were calculated using the R packages `adegenet` (Jombart, 2008) and `hierfstat` (Goudet & Jombart, 2004).

## 3. Results

### 3.1 Genomic predictive ability (PA) and heritability (*h*^2^) estimates

Tables 1 and 2 summarize the median PA and *h*^2^ values for phenotypes estimated using WISER, LS-means, and BLUP across all species and traits, for the GBLUP model. Supplementary materials present median PA values for additional prediction models (random forest, SVR, RKHS, and LASSO), which exhibit, for WISER, LS-means, and BLUP, consistent rankings, similar deviations in median PA values and comparable PA ranges across most models. Therefore, only the GBLUP results are presented here for conciseness. Supplementary Figures 2 to 5 also present a graphical summary of the GBLUP PA values, organized by species, for phenotypes estimated using WISER, LS-means, and BLUP.

**Table 1.** GBLUP median PA values for phenotypic estimation methods (WISER, LS-means, and BLUP) across all species and traits, with interquartile ranges (IQR) in parentheses. Values are based on 100 cross-validation (CV) scenarios.

| Dataset | Trait | GBLUP median PA values |  |  |
| --- | --- | --- | --- | --- |
|  |  | WISER | LS-means | BLUP |
| Rice dataset<br>334 genotypes<br>9,929 SNPs | days to flowering (FL) | 0.74 (0.05) | 0.41 (0.12) | 0.29 (0.15) |
|  | plant height (PH) | 0.73 (0.08) | 0.51 (0.09) | 0.46 (0.10) |
|  | grain yield (YLD) | 0.62 (0.08) | 0.49 (0.11) | 0.42 (0.10) |
|  | grain zinc concentration (ZN) | 0.69 (0.05) | 0.36 (0.16) | 0.32 (0.16) |
| Maize dataset<br>246 genotypes<br>41,723 SNPs | anthesis | 0.74 (0.07) | 0.69 (0.07) | 0.57 (0.10) |
|  | anthesis-silking interval | 0.66 (0.07) | 0.37 (0.18) | 0.22 (0.18) |
|  | ear height | 0.67 (0.07) | 0.24 (0.15) | 0.24 (0.13) |
|  | grain number | 0.68 (0.09) | 0.53 (0.14) | 0.34 (0.16) |
|  | grain weight | 0.61 (0.10) | 0.54 (0.13) | 0.22 (0.16) |
|  | grain yield | 0.77 (0.08) | 0.66 (0.10) | 0.48 (0.14) |
|  | plant height | 0.67 (0.08) | 0.22 (0.15) | 0.22 (0.14) |
|  | silking | 0.70 (0.08) | 0.67 (0.06) | 0.55 (0.08) |
|  | tassel height | 0.67 (0.07) | 0.30 (0.14) | 0.18 (0.15) |
| Apple dataset<br>534 genotypes<br>50,000 SNPs | color over | 0.78 (0.05) | 0.70 (0.05) | 0.56 (0.09) |
|  | flowering begin | 0.92 (0.02) | 0.64 (0.09) | 0.55 (0.09) |
|  | flowering end | 0.84 (0.04) | 0.44 (0.10) | 0.41 (0.10) |
|  | flowering full | 0.80 (0.04) | 0.40 (0.10) | 0.38 (0.10) |
|  | flowering intensity | 0.66 (0.09) | 0.52 (0.09) | 0.32 (0.11) |
|  | fruit number | 0.78 (0.04) | 0.67 (0.06) | 0.36 (0.10) |
|  | fruit weight | 0.78 (0.04) | 0.65 (0.06) | 0.38 (0.10) |
|  | fruit weight single | 0.73 (0.05) | 0.58 (0.07) | 0.54 (0.09) |
|  | harvest date | 0.92 (0.02) | 0.76 (0.06) | 0.59 (0.07) |
|  | powdery mildew | 0.56 (0.08) | 0.40 (0.10) | 0.39 (0.10) |
|  | russet frequency | 0.60 (0.07) | 0.46 (0.09) | 0.36 (0.10) |
|  | scab | 0.56 (0.09) | 0.33 (0.10) | 0.33 (0.10) |
|  | trunk diameter | 0.66 (0.06) | 0.50 (0.06) | 0.36 (0.10) |
|  | trunk increment | 0.51 (0.07) | 0.49 (0.06) | 0.35 (0.10) |
| Pine dataset<br>208 genotypes<br>20,796 SNPs | D | 0.66 (0.10) | 0.08 (0.19) | 0.11 (0.19) |
|  | H | 0.72 (0.08) | 0.18 (0.17) | 0.19 (0.15) |
|  | I | 0.35 (0.25) | 0.10 (0.21) | 0.20 (0.20) |
|  | T4 | 0.34 (0.15) | 0.24 (0.18) | 0.33 (0.21) |
|  | T5 | 0.24 (0.27) | 0.14 (0.22) | 0.18 (0.27) |
|  | T6 | 0.20 (0.27) | 0.14 (0.24) | 0.11 (0.28) |

**Table 2.** GBLUP median *h*^2^ values for phenotypic estimation methods (WISER, LS-means, and BLUP) across all species and traits, with interquartile ranges (IQR) in parentheses. Values are based on 100 cross-validation (CV) scenarios.

| Dataset | Trait | GBLUP median $h^2$ values | | |
| --- | --- | --- | --- | --- |
|  |  | WISER | LS-means | BLUP |
| Rice dataset<br>334 genotypes<br>9,929 SNPs | days to flowering (FL) | 0.87 (0.02) | 0.53 (0.06) | 0.33 (0.11) |
|  | plant height (PH) | 0.86 (0.03) | 0.76 (0.03) | 0.73 (0.04) |
|  | grain yield (YLD) | 0.82 (0.04) | 0.73 (0.05) | 0.68 (0.06) |
|  | grain zinc concentration (ZN) | 0.84 (0.03) | 0.55 (0.10) | 0.50 (0.12) |
| Maize dataset<br>246 genotypes<br>41,723 SNPs | anthesis | 0.96 (0.02) | 0.93 (0.03) | 0.90 (0.03) |
|  | anthesis-silking interval | 0.82 (0.05) | 0.66 (0.11) | 0.55 (0.15) |
|  | ear height | 0.69 (0.05) | 0.60 (0.14) | 0.55 (0.14) |
|  | grain number | 0.96 (0.02) | 0.86 (0.06) | 0.78 (0.09) |
|  | grain weight | 0.65 (0.06) | 0.83 (0.06) | 0.63 (0.16) |
|  | grain yield | 0.97 (0.02) | 0.93 (0.04) | 0.88 (0.07) |
|  | plant height | 0.68 (0.05) | 0.34 (0.15) | 0.33 (0.15) |
|  | silking | 0.71 (0.04) | 0.92 (0.04) | 0.90 (0.05) |
|  | tassel height | 0.68 (0.04) | 0.54 (0.12) | 0.20 (0.38) |
| Apple dataset<br>534 genotypes<br>50,000 SNPs | color over | 0.90 (0.03) | 0.84 (0.03) | 0.80 (0.05) |
|  | flowering begin | 1.00 (0.01) | 0.94 (0.02) | 0.93 (0.03) |
|  | flowering end | 0.96 (0.01) | 0.73 (0.03) | 0.72 (0.04) |
|  | flowering full | 0.94 (0.02) | 0.67 (0.06) | 0.67 (0.05) |
|  | flowering intensity | 0.77 (0.03) | 0.66 (0.04) | 0.56 (0.07) |
|  | fruit number | 0.76 (0.03) | 0.65 (0.04) | 0.47 (0.07) |
|  | fruit weight | 0.75 (0.03) | 0.59 (0.03) | 0.47 (0.05) |
|  | fruit weight single | 0.93 (0.02) | 0.81 (0.04) | 0.79 (0.05) |
|  | harvest date | 1.00 (0.00) | 0.92 (0.02) | 0.88 (0.02) |
|  | powdery mildew | 0.74 (0.04) | 0.51 (0.05) | 0.51 (0.05) |
|  | russet frequency | 0.81 (0.04) | 0.67 (0.06) | 0.54 (0.10) |
|  | scab | 0.68 (0.04) | 0.44 (0.05) | 0.44 (0.04) |
|  | trunk diameter | 0.82 (0.03) | 0.59 (0.04) | 0.52 (0.05) |
|  | trunk increment | 0.59 (0.04) | 0.53 (0.04) | 0.46 (0.06) |
| Pine dataset<br>208 genotypes<br>20,796 SNPs | D | 0.95 (0.03) | 0.00 (0.17) | 0.00 (0.07) |
|  | H | 0.97 (0.01) | 0.30 (0.39) | 0.00 (0.00) |
|  | I | 0.73 (0.16) | 0.00 (0.01) | 0.00 (0.00) |
|  | T4 | 0.63 (0.16) | 0.54 (0.22) | 0.62 (0.22) |
|  | T5 | 0.47 (0.32) | 0.30 (0.44) | 0.26 (0.51) |
|  | T6 | 0.28 (0.34) | 0.08 (0.31) | 0.00 (0.11) |

Table 1 highlights the superior performance of WISER in achieving higher GBLUP median PA values across all species and traits. LS-means generally outperformed BLUP, except in a few isolated cases, while BLUP consistently showed the lowest median PA values. This trend observed for BLUP can be attributed to the assumption of independent and identically distributed (I.I.D.) genetic values in mixed models, combined with imbalanced genotype frequencies, which biases the phenotypic estimation process for each genotype, as discussed in Section 4.1. It is worth noting that, the GBLUP median PA values, for LS-means in Table 1, are consistent with the genomic PA values for the corresponding traits reported in Baertschi et al. (2021), Millet et al. (2019), Jung et al., (2022), and Perry et al. (2022), despite differences in CV schemes and associated scenarios in these studies. This emphasizes the improvement that WISER offers over currently used phenotypic estimation methods.

For the pine dataset, GBLUP median PA values were notably lower, particularly for LS-means and BLUP-estimated phenotypes for traits D, H, and I. These traits exhibited a strong association between genotypes and environments, as indicated by high Cramér’s V statistics (0.43 for D, 1.00 for H, and 0.41 for I). This association resulted in confounding due to the interrelationship between genotypes and environments, leading to perfect multicollinearity among their dummy variables. For example, the variance inflation factor (VIF) for the linear regression used by LS-means was uncomputable for these traits, and similarly, the restricted maximum likelihood (REML) algorithm in BLUP computation failed to converge to identifiable models. In contrast, WISER mitigated these issues through its whitening process, which reduced the confounding relationship between genotypes and fixed-effect variables, particularly the environment. This resulted in more stable and reliable phenotypic estimates. In terms of interquartile ranges (IQR) for GBLUP PA values, WISER typically produced smaller ranges compared to LS-means and BLUP, with a few exceptions, most notably in the pine dataset. Across all species and traits, the average IQR values were 0.09 for WISER, 0.12 for LS-means, and 0.13 for BLUP, highlighting WISER’s greater robustness in most scenarios.

To evaluate the potential improvements WISER offers over other existing phenotypic estimation methods, and considering LS-means’ superior performance compared to BLUP, only the deviations between the GBLUP median PA values of WISER and LS-means—referred to as the median PA gain— were calculated. The average median PA gain across all species and traits was 0.22, demonstrating WISER’s enhanced accuracy in phenotypic estimation.

Table 2 demonstrates similar trends for GBLUP median *h*^2^ values, with WISER consistently outperforming LS-means and BLUP across all species and traits, except for two isolated cases: grain weight and silking. However, it is important to note that a higher *h*^2^ estimate does not always indicate a better estimation, as it may result from overfitting of the GBLUP model, which can lead to an overestimation of the trait’s heritability. For instance, while the *h*^2^ values estimated using LS-means phenotypes were higher than those obtained with WISER for grain weight and silking, the corresponding GBLUP median PA values based on LS-means phenotypes were lower than those derived from WISER phenotypes. For traits D, H, and I in the pine dataset, LS-means and BLUP exhibited exceptionally low *h*^2^ values due to numerical instability caused by the strong confounding associations between genotypes and environments. These values, rounded to two decimal places, are not actual zeros but rather reflect challenges in model convergence.

### 3.2. Factors contributing to the increase in median PA gain

To further investigate the factors contributing to the increase in median PA gain, correlations were computed between the median PA gain and several statistics defined in Section 2.2: deviations in median *h*^2^ values between WISER and LS-means, Shannon entropy of genotype frequencies, Shannon joint entropy of genotype and site (or environment), and the *F*_*ST*_ calculated from the clusters estimated through the combination of UMAP and *K*-means applied to SNP marker data. These analyses provide valuable insights into the underlying mechanisms driving WISER’s improved performance, particularly its capacity to address confounding relationships. Note that PCA was not used for dimensionality reduction prior to *K*-means, as it failed to distinguish known accessions and 27 progeny families in the apple REFPOP dataset (see Supplementary Figures 6 to 9). In contrast, UMAP, without any labeling information, was able to effectively separate the accessions and most of the families, making it better suited for use prior to the clustering algorithm.

Figure 1 presents correlation plots between the median PA gain values and various statistics, as defined in Section 2.3.4, for the 33 traits across species. A strong positive correlation of 0.70 was observed between the median PA gain values and the median *h*^2^ deviations between WISER and LS-means phenotypes for these traits. This result aligns with expectations, as an increase in *h*^2^ estimates typically reflects higher genetic variance estimates and better predictive ability. In contrast, weak to moderate negative correlations were found between the median PA gain values and the Shannon entropy of genotype frequencies (−0.22), as well as the Shannon joint entropies of genotype and site (−0.34) and genotype and environment (−0.38). Although these correlations were not strong, they suggested a trend where the median PA gain achieved by WISER tended to increase in scenarios with higher genotype-site or genotype-environment confounding.

**Figure 1.**
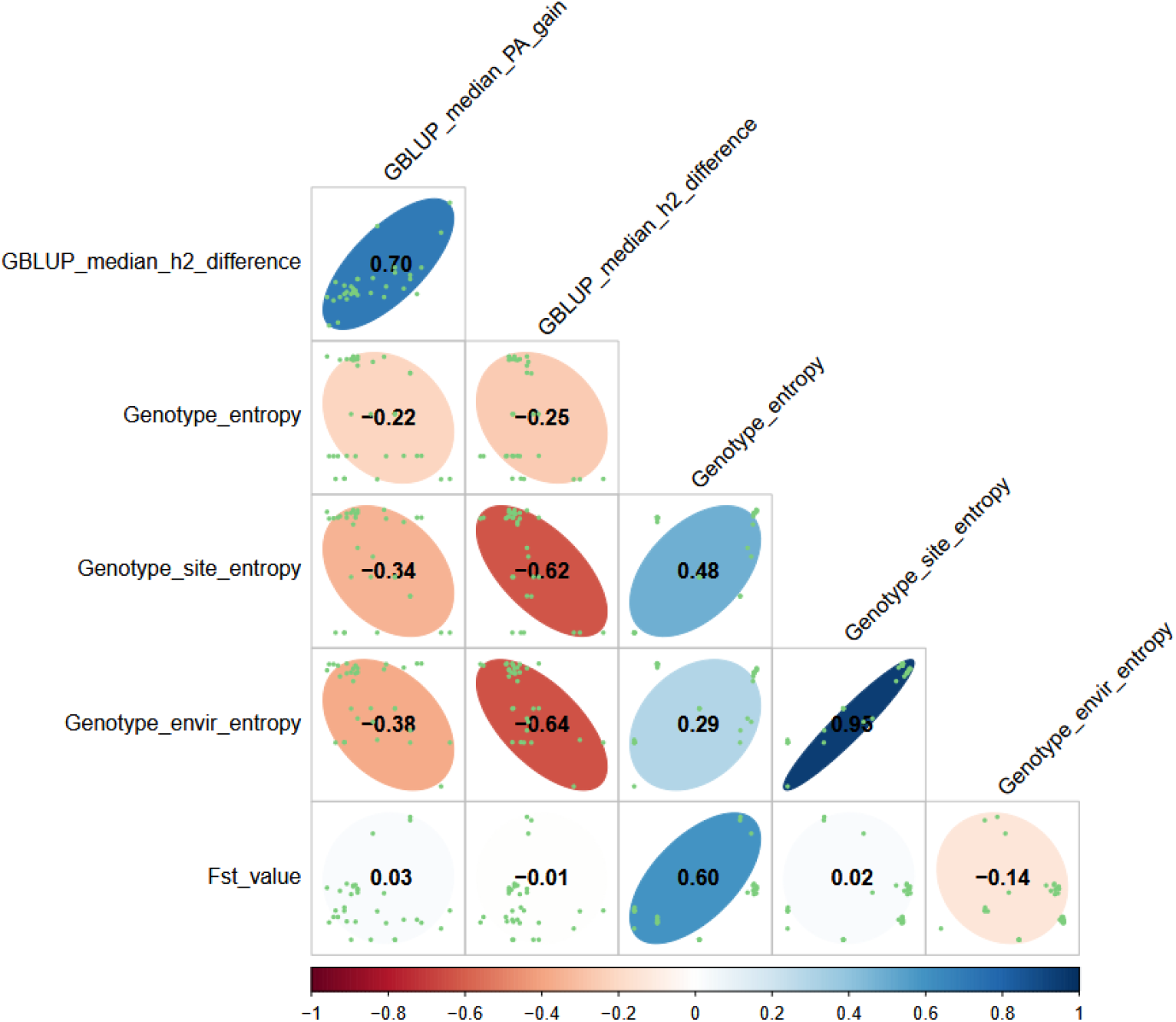
Correlation plots between GBLUP median PA gain, median *h*^2^ deviations, Shannon entropies, and *F*_*ST*_ values across all species and traits.

Notably, when considering the median *h*^2^ deviations rather than the median PA gain, stronger trends emerged. More pronounced negative correlations with the Shannon joint entropies of genotype and site (−0.62) and genotype and environment (−0.64) were observed, highlighting the significant impact of the confounding between genotype and site or environment on genetic variance estimation. On the other hand, correlations close to zero were observed between the median PA gain and *F*_*ST*_, as well as between the median *h*^2^ deviations and *F*_*ST*_, making the relationship inconclusive.

The weak negative correlations, and those close to zero, observed in Figure 1 are likely unreliable and attributable to various species-specific factors, such as marker density, linkage disequilibrium patterns, and allele and genotype frequencies, which exhibit minimal or no variation between traits within each species. Table 3 presents the median values and interquartile ranges (IQR) of Shannon entropy for genotype frequencies (*H*(*Genotype*)) and *F*_*ST*_ values across traits for each species. Notably, it shows limited or no variation in these statistics among traits within each species, which likely contributed to the reduced reliability of the correlations.

**Table 3.** Median and IQR values for *H*(*Genotype*) and *F*_*ST*_ across traits per species.

| Species | $median(H(Genotype))$ | $IQR(H(Genotype))$ | $median(F_{ST})$ | $IQR(F_{ST})$ |
| --- | --- | --- | --- | --- |
| Rice | 8.365 | 0.000 | 0.010 | 0.000 |
| Maize | 7.932 | 0.001 | 0.026 | 0.002 |
| Apple | 8.929 | 0.042 | 0.056 | 0.006 |
| Pine | 7.695 | 0.005 | 0.035 | 0.001 |

Correlation plots similar to those in Figure 1 were generated for the apple dataset, which contained the highest number of traits (14) among all datasets. This enabled an exploration of intra-species relationships in at least one species (see Supplementary Figure 10). However, the interpretation of these plots was not possible due to the small number of traits, as meaningful calculated correlations typically require a larger sample size. Nevertheless, the trends observed in these plots were consistent with those seen in the inter-species analysis, particularly for the correlations between the median PA gain values, the median *h*^2^ deviations, and the Shannon entropies. Additionally, a positive correlation between the median PA gain and *F*_*ST*_ was observed, but this correlation is likely coincidental due to the small number of points, their limited variability, and observed dispersion in the corresponding plot.

In summary, the negative correlations observed between the median PA gain values (or the median *h*^2^ deviations) and the Shannon entropies highlight WISER’s ability to improve phenotypic estimation in the presence of confounding factors and population complexity.

## 4. Discussion

This study demonstrates the significant improvement in phenotypic estimation offered by the WISER approach compared to traditional methods like LS-means and BLUP, across multiple datasets and traits. The results presented here highlight WISER’s ability to handle genotype-environment confounding relationships, genotype frequency imbalances, and complex population structures, which are major challenges in omics-based selection and association studies.

### 4.1 Comparison of WISER, LS-means, and BLUP

The findings demonstrate that WISER consistently outperforms both LS-means and BLUP in terms of median PA and *h*^2^ estimates. Across all species and traits, WISER achieved the highest GBLUP median PA values, followed by LS-means and BLUP, which often showed the lowest accuracy. This trend aligns with the known limitations of BLUP, particularly in scenarios with imbalanced genotype frequencies and the assumption of I.I.D. genetic values, which can lead to biased phenotypic estimates (Holland & Piepho, 2024).

Within the WISER framework, the vector ν of phenotypes is not assumed to be a random vector drawn from a distribution with a specified covariance matrix. This design avoids imposing an unnecessary covariance structure during the estimation of ν, which could lead to undesirable consequences. For example, assuming 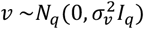 is often unrealistic and results in a decorrelated covariance structure in the best linear unbiased predictor (BLUP) of ν. This is particularly problematic due to BLUP’s inherent properties as a shrinkage estimator, especially in the context of imbalanced datasets. For instance, Holland & Piepho (2024) emphasize that BLUP estimates, when used as phenotypes, are not recommended if they are derived under the assumption that genetic values are I.I.D. when the data are unbalanced—a very frequent situation due to missing data or incomplete block experimental designs. In such situations, the combination of the I.I.D. assumption and data imbalance leads to shrinkage of genotype estimates toward the population mean, with the degree of shrinkage being inversely proportional to the number of replicates for each genotype. This effect is particularly problematic when many genotypes have few replicates, as their BLUP values will disproportionately shrink toward the mean, distorting genetic rankings and undermining the interpretability and utility of these estimates for subsequent analyses.

The successive ordinary least squares (OLS) procedure implemented in WISER addresses these issues by operating without such assumptions. It ensures that the estimation of ν is unrelated to omic information and pre-defined covariance structures, treating ν as a distinct vector of fixed effects. This approach aligns with the recommendations of Holland & Piepho (2024), providing reliable phenotypic estimates and preserving genetic value rankings, even in unbalanced experimental designs. Moreover, by maintaining the separation of ν from omic information, the approach preserves the genetic signals essential for association analyses, ensuring that no valuable information is inadvertently filtered out—an issue that would arise if phenotypes were estimated using a polygenic background based on omic data.

While LS-means was generally more robust than BLUP, it still underperformed compared to WISER, especially in datasets where genotype-environment associations were strong and multicollinearity was severe. WISER’s superior performance can be attributed to its whitening process, which reduces the confounding relationships between genotypes and fixed-effect variables, such as environment. By mitigating such confounding relationships, WISER yields more reliable and stable estimates, especially for datasets where traditional approaches for phenotypic estimation struggle with multicollinearity, as seen in the pine dataset for traits with strong genotype-environment associations.

### 4.2. Impact of confounding effects and imbalanced frequencies

The study also explored how confounding effects and imbalanced frequencies influence the accuracy of phenotypic estimation. WISER was found to be particularly effective in mitigating these challenges. For instance, in the pine dataset, where traits such as D, H, and I exhibited strong genotype-environment associations and confounding, WISER provided more reliable phenotypic estimates compared to LS-means and BLUP. This was because the REML algorithm in BLUP and the linear regression (prior to LS-means) encountered issues such as non-optimal convergence and multicollinearity, respectively.

This finding is critical, as genotype-environment confounding relationships, often exacerbated by high variability in genotype frequencies across environments, can severely compromise the accuracy of phenotypic estimations and predictions. Moreover, imbalances in genotype frequencies, along with their variability across environments, inevitably affect the genetic covariance structure in experimental designs. This underscores the complex interdependence of these factors and their combined impact on phenotypic estimation. WISER’s whitening process addresses these issues by simultaneously mitigating confounding effects, multicollinearity, and imbalances in genotype frequencies, ultimately improving the accuracy and robustness of phenotypic estimates.

Notably, some degree of confounding, imbalance in genotype frequencies, and population structure are nearly inevitable, even in carefully balanced experimental designs. This is exemplified by the apple REFPOP dataset, which, despite being explicitly designed for balance, still displayed residual imbalances and confounding relationships. This highlights the pervasive nature of these challenges in real-world datasets and underscores the importance of methods like WISER, which can effectively address such complexities.

### 4.3. Impact of the degree of population structure

The study further explored the relationship between improvements in phenotypic estimation for each trait using WISER and the degree of population structure, quantified through genetic differentiation. However, the relationship remained inconclusive due to species-specific factors and limited to no variation in the genomic data across traits for each species. Nonetheless, WISER proved effective in removing the confounding effects of population structure on fixed-effect variables, particularly environmental effects. Among the various computed statistics, the strongest negative correlations were observed between both the median PA gain and the median *h*^2^ deviations, and the Shannon joint entropy of genotype and environment.

Hence, these findings point out the critical importance of considering population structure when selecting methodologies for phenotypic estimation. As demonstrated in this study, population structure significantly impacts genomic predictive ability through confounding effects between genotypes and environments, and it is a well-established factor influencing association studies (Azevedo et al., 2017; Daetwyler et al., 2012; Kang et al., 2008; Patterson et al., 2006; Price et al., 2006; Riedelsheimer et al., 2012; Yu et al., 2006).

### 4.4. Phenotypic estimation, genetic variance, and predictive ability

The positive correlation between the median PA gain and the median *h*^2^ deviations underscores the relationships between phenotypic estimation, genetic variance, and predictive ability. Across all species and traits, a strong positive correlation (0.70) was observed between the median PA gain and the median *h*^2^ deviations. This demonstrates that WISER-estimated phenotypes improve the capture of genetic variance when using GBLUP, thereby enhancing predictions. Interestingly, the median *h*^2^ deviations exhibited strong negative correlations with the Shannon joint entropies of genotype and site or environment. Similarly, moderate negative correlations were observed between the median PA gain values and these Shannon joint entropies. These trends were also observed in the apple dataset, despite the low number of traits in this dataset limiting the interpretability. These findings suggest that WISER is particularly effective in addressing scenarios characterized by a high degree of genotype-site and/or genotype-environment confounding, thereby improving phenotypic estimation in these complex situations. This emphasizes WISER’s ability to deliver accurate phenotypic estimates in the presence of such confounding effects, while also improving genetic variance estimation and predictive ability.

### 4.5. Limitations, computational efficiency, and population size considerations

WISER’s strengths are most apparent when multiple fixed-effect variables are considered, particularly in defining the environmental variable. However, its utility is limited in experimental designs with very few fixed-effect variables (e.g., only an overall mean and one additional factor), as these do not permit a more precise definition of the environmental variable. This highlights the importance of comprehensive experimental designs to fully harness WISER’s potential. While computational efficiency may have been a concern in the past, the current availability of high-performance computing (HPC) clusters has largely mitigated this issue. For example, processing times for the rice and pine datasets were under five hours, depending on resource availability and user demand on the HPC cluster. Furthermore, WISER employs parallelized algorithms, such as an ABC approach, which ensures rapid and stable variance component estimation, even for large datasets linked to complex experimental designs. This approach is preferable to iterative algorithms, such as expectation-maximization (EM), which can be slow, produce inaccurate results, or converge to suboptimal solutions, especially when handling large datasets (Bradley et al., 1998; Ordonez & Omiecinski, 2002; L. Xu & Jordan, 1996). Nonetheless, it is important to note that WISER may encounter scalability challenges when applied to datasets with populations significantly larger than those in the apple REFPOP dataset. Future work should explore WISER’s performance on larger datasets.

### 4.5. Historical context, implications for breeding programs, versatility of the whitening approach and future directions

Historically, tools for managing confounding effects due to population structure and imbalanced genotype frequencies in phenotypic estimation were limited. This led to the widespread use of methods like BLUP. BLUP remains a popular choice due to its simplicity and the availability of user-friendly R packages like `lme4` (Bates et al., 2015). However, BLUP’s reliance on the I.I.D. assumption in this usage context and its sensitivity to genotype frequency imbalances result in known suboptimal performance, as pointed out by Holland & Piepho (2024). The `wiser` R package addresses this gap by offering a more robust framework for phenotypic estimation, specifically tailored for experimental setups with population structure, confounding relationships, and imbalances in genotype frequencies. By overcoming these challenges, WISER has significant implications for breeding and genetic selection programs. Accurate phenotypic estimation is critical for predicting complex traits and selecting individuals with desirable genetic profiles. The reduced interquartile ranges (IQR) for GBLUP PA values observed with WISER demonstrate its ability to deliver more consistent predictions across cross-validation scenarios, a feature invaluable for breeders seeking to improve the reliability of selection decisions. Furthermore, the whitening approach employed by WISER has broader applications beyond omic data. For instance, in image analysis, whitening techniques are effectively used to preprocess non-random vectors, such as images (Ermolov et al., 2021; Gluckman, 2005). This underscores the versatility of the whitening approach, demonstrating its adaptability to diverse types of data. Looking ahead, the capacity of WISER to handle more complex datasets with additional environmental covariates represents an exciting avenue for further research. Future studies could also explore integrating WISER with advanced predictive modeling techniques, such as deep learning, to enhance phenotypic predictive accuracy. Furthermore, incorporating genotype-environment interactions into WISER’s estimation process represents a promising direction for refining phenotypic estimates. These advancements have the potential to unlock even greater possibilities for phenotypic estimation and omic-based selection, driving innovation in breeding programs.

## 5. Conclusion

In conclusion, the results of this study provide strong evidence that WISER offers significant advantages in phenotypic estimation, particularly in datasets with complex genotype-environment confounding relationships, genotype frequency imbalances, and population structure. By addressing these challenges, WISER enables more accurate and stable phenotypic estimations, and predictions, which could lead to improved genetic selection strategies in crop and livestock breeding programs.

## Author contributions

LJ derived the algebra for the analytical results, created the GitHub repositories, developed the associated R scripts, conducted all dataset analyses, and developed the WISER R package. LJ also authored the manuscript. WG, ML, AP, MR, CD, FL, MA, and HM reviewed and approved the final version of the manuscript.

## Data availability

All datasets and scripts used in this study are publicly available on GitHub:

- https://github.com/ljacquin/wiser_genomic_prediction_rice
- https://github.com/ljacquin/wiser_genomic_prediction_maize
- https://github.com/ljacquin/refpop
- https://github.com/ljacquin/wiser_genomic_prediction_pine
- https://github.com/ljacquin/compute_stats_wiser_results

## Acknowledgments

The authors thank the field technicians and staff at INRAe IRHS and Experimental Unit (UE Horti), Angers, France, the Fruit Breeding Group at Agroscope in Waedenswil, Switzerland, and technical staff at all apple REFPOP sites for the maintenance of the orchards and phenotypic data collection. Phenotypic data collection was partially supported by the Horizon 2020 Framework Program of the European Union under grant agreement No 817970 (project INVITE: “Innovations in plant variety testing in Europe to foster the introduction of new varieties better adapted to varying biotic and abiotic conditions and to more sustainable crop management practices”) and by the SusCrop Agrobiodiversity project Apple BIOME (“Microbiome and genomic analysis in apple germplasm towards broadening genetic resources to breed for resilient varieties”). LJ was supported by the ANR grant for the Apple BIOME project (ANR-22-SUSC-0001-05).

## Literature cited

Amadeu, R. R., Ferrão, L. F. V., Oliveira, I. de B., Benevenuto, J., Endelman, J. B., & Munoz, P. R. (2020). Impact of dominance effects on autotetraploid genomic prediction. Crop Science, 60(2), 656–665. 10.1002/csc2.20075

Astle, W., & Balding, D. J. (2009). Population Structure and Cryptic Relatedness in Genetic Association Studies. Statistical Science, 24(4), 451–471. 10.1214/09-STS307

Azevedo, C. F., Resende M. D. V. de, Silva, F. F. e, Nascimento, M., Viana, J. M. S., & Valente, M. S. F. (2017). Population structure correction for genomic selection through eigenvector covariates. Crop Breeding and Applied Biotechnology, 17, 350–358. 10.1590/1984-70332017v17n4a53

Baertschi, C., Cao, T.-V., Bartholomé, J., Ospina, Y., Quintero, C., Frouin, J., Bouvet, J.-M., & Grenier, C. (2021). Impact of early genomic prediction for recurrent selection in an upland rice synthetic population. G3 Genes|Genomes|Genetics, 11(12), jkab320. 10.1093/g3journal/jkab320

Bates, D., Mächler, M., Bolker, B., & Walker, S. (2015). Fitting Linear Mixed-Effects Models Using lme4. Journal of Statistical Software, 67, 1–48. 10.18637/jss.v067.i01

Beugin, M.-P., Gayet, T., Pontier, D., Devillard, S., & Jombart, T. (2018). A fast likelihood solution to the genetic clustering problem. Methods in Ecology and Evolution, 9(4), 1006–1016. 10.1111/2041-210X.12968

Bradley, P., Fayyad, U., & Reina, C. (1998, novembre 1). Scaling EM (Expectation Maximization) Clustering to Large Databases. https://www.semanticscholar.org/paper/Scaling-EM-(Expectation-Maximization)-Clustering-to-Bradley-Fayyad/80186e214b942304e6dfba8039ae1a16ee3ec270

Byrne, R. P., van Rheenen, W., van den Berg, L. H., Veldink, J. H., & McLaughlin, R. L. (2020). Dutch population structure across space, time and GWAS design. Nature Communications, 11(1), 4556. 10.1038/s41467-020-18418-4

Carress, H., Lawson, D. J., & Elhaik, E. (2021). Population genetic considerations for using biobanks as international resources in the pandemic era and beyond. BMC Genomics, 22(1), 351. 10.1186/s12864-021-07618-x

Chen, Y., Gao, Y., Chen, P., Zhou, J., Zhang, C., Song, Z., Huo, X., Du, Z., Gong, J., Zhao, C., Wang, S., Zhang, J., Wang, F., & Zhang, J. (2022). Genome-wide association study reveals novel quantitative trait loci and candidate genes of lint percentage in upland cotton based on the CottonSNP80K array. Theoretical and Applied Genetics, 135(7), 2279–2295. 10.1007/s00122-022-04111-1

Cherkassky, V., & Ma, Y. (2004). Practical selection of SVM parameters and noise estimation for SVM regression. Neural Networks, 17(1), 113–126. 10.1016/S0893-6080(03)00169-2

Daetwyler, H. D., Kemper, K. E., van der Werf, J. H. J., & Hayes, B. J. (2012). Components of the accuracy of genomic prediction in a multi-breed sheep population. Journal of Animal Science, 90(10), 3375–3384. 10.2527/jas.2011-4557

Elhaik, E. (2022). Principal Component Analyses (PCA)-based findings in population genetic studies are highly biased and must be reevaluated. Scientific Reports, 12(1), 14683. 10.1038/s41598-022-14395-4

Ermolov, A., Siarohin, A., Sangineto, E., & Sebe, N. (2021). Whitening for Self-Supervised Representation Learning. Proceedings of the 38th International Conference on Machine Learning, 3015–3024. https://proceedings.mlr.press/v139/ermolov21a.html

Friedman, J., Hastie, T., & Tibshirani, R. (2010). Regularization Paths for Generalized Linear Models via Coordinate Descent. Journal of statistical software, 33(1), 1–22.

Gloss, A. D., Steiner, M. C., Novembre, J., & Bergelson, J. (2023). The design of mapping populations: Impacts of geographic scale on genetic architecture and mapping efficacy for defense and immunity. Current Opinion in Plant Biology, 74, 102399. 10.1016/j.pbi.2023.102399

Gluckman, J. (2005). Higher order whitening of natural images. 2005 IEEE Computer Society Conference on Computer Vision and Pattern Recognition (CVPR’05), 2, 354–360 vol. 2. 10.1109/CVPR.2005.175

Goudet, J., & Jombart, T. (2004). hierfstat: Estimation and Tests of Hierarchical F-Statistics (p. 0. 5–11) [Jeu de données]. 10.32614/CRAN.package.hierfstat

Henderson, C. R. (1984). Applications of linear models in animal breeding. University of Guelph.

Hennig, C. (2003). fpc: Flexible Procedures for Clustering (p. 2. 2–13) [Jeu de données]. 10.32614/CRAN.package.fpc

Hoffman, G. E. (2013). Correcting for Population Structure and Kinship Using the Linear Mixed Model: Theory and Extensions. PLOS ONE, 8(10), e75707. 10.1371/journal.pone.0075707

Holland, J. B., & Piepho, H.-P. (2024). Don’t BLUP Twice. G3 Genes|Genomes|Genetics, 14(12), jkae250. 10.1093/g3journal/jkae250

Hu, H., Campbell, M. T., Yeats, T. H., Zheng, X., Runcie, D. E., Covarrubias-Pazaran, G., Broeckling, C., Yao, L., Caffe-Treml, M., Gutiérrez, L., Smith, K. P., Tanaka, J., Hoekenga, O. A., Sorrells, M. E., Gore, M. A., & Jannink, J.-L. (2021). Multi-omics prediction of oat agronomic and seed nutritional traits across environments and in distantly related populations. Theoretical and Applied Genetics, 134(12), 4043–4054. 10.1007/s00122-021-03946-4

Jacquin, L., Cao, T.-V., & Ahmadi, N. (2016). A Unified and Comprehensible View of Parametric and Kernel Methods for Genomic Prediction with Application to Rice. Frontiers in Genetics, 7. 10.3389/fgene.2016.00145

Jombart, T. (2008). adegenet: A R package for the multivariate analysis of genetic markers. Bioinformatics (Oxford, England), 24(11), 1403–1405. 10.1093/bioinformatics/btn129

Jung, M., Keller, B., Roth, M., Aranzana, M. J., Auwerkerken, A., Guerra, W., Al-Rifaï, M., Lewandowski, M., Sanin, N., Rymenants, M., Didelot, F., Dujak, C., Font I Forcada, C., Knauf, A., Laurens, F., Studer, B., Muranty, H., & Patocchi, A. (2022). Genetic architecture and genomic predictive ability of apple quantitative traits across environments. Horticulture Research, 9, uhac028. 10.1093/hr/uhac028

Jung, M., Roth, M., Aranzana, M. J., Auwerkerken, A., Bink, M., Denancé, C., Dujak, C., Durel, C.-E., Font I Forcada, C., Cantin, C. M., Guerra, W., Howard, N. P., Keller, B., Lewandowski, M., Ordidge, M., Rymenants, M., Sanin, N., Studer, B., Zurawicz, E., … Muranty, H. (2020). The apple REFPOP—a reference population for genomics-assisted breeding in apple. Horticulture Research, 7(1), 1–16. 10.1038/s41438-020-00408-8

Kang, H. M., Zaitlen, N. A., Wade, C. M., Kirby, A., Heckerman, D., Daly, M. J., & Eskin, E. (2008). Efficient control of population structure in model organism association mapping. Genetics, 178(3), 1709–1723. 10.1534/genetics.107.080101

Karatzoglou, A., Smola, A., Hornik, K., & Zeileis, A. (2004). kernlab—An S4 Package for Kernel Methods in R. Journal of Statistical Software, 11, 1–20. 10.18637/jss.v011.i09

Kessy, A., Lewin, A., & Strimmer, K. (2018). Optimal Whitening and Decorrelation. The American Statistician, 72(4), 309–314. 10.1080/00031305.2016.1277159

Konopka, T. (2023). umap: Uniform Manifold Approximation and Projection (Version 0.2.10.0) [Logiciel]. https://cran.r-project.org/web/packages/umap/index.html

Korbinian Strimmer, Takoua Jendoubi, Agnan Kessy, Alex Lewin. (2018). whitening: Whitening and High-Dimensional Canonical Correlation Analysis (p. 1.4.0) [Jeu de données]. 10.32614/CRAN.package.whitening

Kwan, C. C. Y. (2011). An Introduction to Shrinkage Estimation of the Covariance Matrix: A Pedagogic Illustration. Spreadsheets in Education. https://www.semanticscholar.org/paper/An-Introduction-to-Shrinkage-Estimation-of-the-A-Kwan/a5ad09a33a956405bdc5f829b4e83966c99183f3

Lawson, D. J., Davies, N. M., Haworth, S., Ashraf, B., Howe, L., Crawford, A., Hemani, G., Davey Smith, G., & Timpson, N. J. (2020). Is population structure in the genetic biobank era irrelevant, a challenge, or an opportunity? Human Genetics, 139(1), 23–41. 10.1007/s00439-019-02014-8

Lenth, R. V. (2016). Least-Squares Means: The R Package lsmeans. Journal of Statistical Software, 69, 1–33. 10.18637/jss.v069.i01

Li, H., Cheng, X., Zhang, L., Hu, J., Zhang, F., Chen, B., Xu, K., Gao, G., Li, H., Li, L., Huang, Q., Li, Z., Yan, G., & Wu, X. (2018). An Integration of Genome-Wide Association Study and Gene Co-expression Network Analysis Identifies Candidate Genes of Stem Lodging-Related Traits in Brassica napus. Frontiers in Plant Science, 9. 10.3389/fpls.2018.00796

Liu, L., Zhang, D., Liu, H., & Arendt, C. (2013). Robust methods for population stratification in genome wide association studies. BMC Bioinformatics, 14(1), 132. 10.1186/1471-2105-14-132

Lorenzi, A., Bauland, C., Mary-Huard, T., Pin, S., Palaffre, C., Guillaume, C., Lehermeier, C., Charcosset, A., & Moreau, L. (2022). Genomic prediction of hybrid performance: Comparison of the efficiency of factorial and tester designs used as training sets in a multiparental connected reciprocal design for maize silage. Theoretical and Applied Genetics, 135(9), 3143–3160. 10.1007/s00122-022-04176-y

Lorenzi, A., Bauland, C., Pin, S., Madur, D., Combes, V., Palaffre, C., Guillaume, C., Touzy, G., Mary-Huard, T., Charcosset, A., & Moreau, L. (2024). Portability of genomic predictions trained on sparse factorial designs across two maize silage breeding cycles. Theoretical and Applied Genetics, 137(3), 75. 10.1007/s00122-024-04566-4

McInnes, L., Healy, J., Saul, N., & Großberger, L. (2018). UMAP: Uniform Manifold Approximation and Projection. Journal of Open Source Software, 3(29), 861. 10.21105/joss.00861

McLeod, L., Barchi, L., Tumino, G., Tripodi, P., Salinier, J., Gros, C., Boyaci, H. F., Ozalp, R., Borovsky, Y., Schafleitner, R., Barchenger, D., Finkers, R., Brouwer, M., Stein, N., Rabanus-Wallace, M. T., Giuliano, G., Voorrips, R., Paran, I., & Lefebvre, V. (2023). Multi-environment association study highlights candidate genes for robust agronomic quantitative trait loci in a novel worldwide Capsicum core collection. The Plant Journal, 116(5), 1508–1528. 10.1111/tpj.16425

Meyer, D., Zeileis, A., Hornik, K., & Friendly, M. (2002). vcd: Visualizing Categorical Data (p. 1. 4–13) [Jeu de données]. 10.32614/CRAN.package.vcd

Millet, E. J., Kruijer, W., Coupel-Ledru, A., Alvarez Prado, S., Cabrera-Bosquet, L., Lacube, S., Charcosset, A., Welcker, C., van Eeuwijk, F., & Tardieu, F. (2019). Genomic prediction of maize yield across European environmental conditions. Nature Genetics, 51(6), 952–956. 10.1038/s41588-019-0414-y

Nikovski, D., & Byadarhaly, K. (2016). Regularized covariance matrix estimation with high dimensional data for supervised anomaly detection problems. 2016 International Joint Conference on Neural Networks (IJCNN), 2811–2818. 10.1109/IJCNN.2016.7727554

Njuguna, J. N., Clark, L. V., Lipka, A. E., Anzoua, K. G., Bagmet, L., Chebukin, P., Dwiyanti, M. S., Dzyubenko, E., Dzyubenko, N., Ghimire, B. K., Jin, X., Johnson, D. A., Nagano, H., Peng, J., Petersen, K. K., Sabitov, A., Seong, E. S., Yamada, T., Yoo, J. H., … Sacks, E. J. (2023). Genome-wide association and genomic prediction for yield and component traits of Miscanthus sacchariflorus. GCB Bioenergy, 15(11), 1355–1372. 10.1111/gcbb.13097

Ordonez, C., & Omiecinski, E. (2002). FREM: Fast and robust EM clustering for large data sets. Proceedings of the Eleventh International Conference on Information and Knowledge Management, 590–599. 10.1145/584792.584889

Patterson, N., Price, A. L., & Reich, D. (2006). Population Structure and Eigenanalysis. PLOS Genetics, 2(12), e190. 10.1371/journal.pgen.0020190

Perry, A., Wachowiak, W., Beaton, J., Iason, G., Cottrell, J., & Cavers, S. (2022). Identifying and testing marker–trait associations for growth and phenology in three pine species: Implications for genomic prediction. Evolutionary Applications, 15(2), 330–348. 10.1111/eva.13345

Podosinnikova, A. (2016). On the method of moments for estimation in latent linear models [Phdthesis, Université Paris sciences et lettres]. https://theses.hal.science/tel-01489260

Price, A. L., Patterson, N. J., Plenge, R. M., Weinblatt, M. E., Shadick, N. A., & Reich, D. (2006). Principal components analysis corrects for stratification in genome-wide association studies. Nature Genetics, 38(8), 904–909. 10.1038/ng1847

Pritchard, J. K., Stephens, M., Rosenberg, N. A., & Donnelly, P. (2000). Association Mapping in Structured Populations. The American Journal of Human Genetics, 67(1), 170–181. 10.1086/302959

Riedelsheimer, C., Technow, F., & Melchinger, A. E. (2012). Comparison of whole-genome prediction models for traits with contrasting genetic architecture in a diversity panel of maize inbred lines. BMC Genomics, 13(1), 452. 10.1186/1471-2164-13-452

Rio, S., Gallego-Sánchez, L., Montilla-Bascón, G., Canales, F. J., Isidro Y Sánchez, J., & Prats, E. (2021). Genomic prediction and training set optimization in a structured Mediterranean oat population. TAG. Theoretical and Applied Genetics. Theoretische Und Angewandte Genetik, 134(11), 3595–3609. 10.1007/s00122-021-03916-w

Rodríguez-Álvarez, M. X., Boer, M. P., van Eeuwijk, F. A., & Eilers, P. H. C. (2018). Correcting for spatial heterogeneity in plant breeding experiments with P-splines. Spatial Statistics, 23, 52–71. 10.1016/j.spasta.2017.10.003

Rohart, F., Gautier, B., Singh, A., & Cao, K.-A. L. (2017). mixOmics: An R package for ‘omics feature selection and multiple data integration. PLOS Computational Biology, 13(11), e1005752. 10.1371/journal.pcbi.1005752

Sakhale, S. A., Yadav, S., Clark, L. V., Lipka, A. E., Kumar, A., & Sacks, E. J. (2023). Genome-wide association analysis for emergence of deeply sown rice (Oryza sativa) reveals novel aus-specific phytohormone candidate genes for adaptation to dry-direct seeding in the field. Frontiers in Plant Science, 14, 1172816. 10.3389/fpls.2023.1172816

Samira, R., Kimball, J. A., Samayoa, L. F., Holland, J. B., Jamann, T. M., Brown, P. J., Stacey, G., & Balint-Kurti, P. J. (2020). Genome-wide association analysis of the strength of the MAMP-elicited defense response and resistance to target leaf spot in sorghum. Scientific Reports, 10, 20817. 10.1038/s41598-020-77684-w

Schäfer, J., & Strimmer, K. (2005). A shrinkage approach to large-scale covariance matrix estimation and implications for functional genomics. Statistical Applications in Genetics and Molecular Biology, 4, Article32. 10.2202/1544-6115.1175

Steland, A. (2018). Shrinkage for covariance estimation: Asymptotics, confidence intervals, bounds and applications in sensor monitoring and finance. Statistical Papers, 59(4), 1441–1462. 10.1007/s00362-018-1040-y

Sul, J. H., Martin, L. S., & Eskin, E. (2018). Population structure in genetic studies: Confounding factors and mixed models. PLOS Genetics, 14(12), e1007309. 10.1371/journal.pgen.1007309

Theiler, J. (2012). The incredible shrinking covariance estimator. 83910P-83910P - 12. 10.1117/12.918718

Touloumis, A. (2015). Nonparametric Stein-type shrinkage covariance matrix estimators in high-dimensional settings. Computational Statistics & Data Analysis, 83, 251–261. 10.1016/j.csda.2014.10.018

Tucker, J. R., Brûlé-Babel, A. L., Hiebert, C. W., Larios, R., Legge, W. G., Badea, A., & Fernando, W. G. D. (2022). Genetic structure and genome-wide association study of a genomic panel of two-row, spring barley (Hordeum vulgare L.) with differential reaction to Fusarium head blight (Fusarium graminearum Schwabe) and deoxynivalenol production. Canadian Journal of Plant Pathology, 44(6), 874–891. 10.1080/07060661.2022.2086925

VanRaden, P. M. (2008). Efficient Methods to Compute Genomic Predictions. Journal of Dairy Science, 91(11), 4414–4423. 10.3168/jds.2007-0980

Vogel, G., Giles, G., Robbins, K. R., Gore, M. A., & Smart, C. D. (2022). Quantitative Genetic Analysis of Interactions in the Pepper-Phytophthora capsici Pathosystem. Molecular Plant-Microbe Interactions: MPMI, 35(11), 1018–1033. 10.1094/MPMI-12-21-0307-R

Wang, C., Tindemans, S. H., & Palensky, P. (2024). Improved Anomaly Detection and Localization Using Whitening-Enhanced Autoencoders. IEEE Transactions on Industrial Informatics, 20(1), 659–668. 10.1109/TII.2023.3268685

Wang, M., Fang, Z., Yoo, B., Bejerano, G., & Peltz, G. (2021). The Effect of Population Structure on Murine Genome-Wide Association Studies. Frontiers in Genetics, 12. 10.3389/fgene.2021.745361

Wright, M. N., & Ziegler, A. (2017). ranger: A Fast Implementation of Random Forests for High Dimensional Data in C++ and R. Journal of Statistical Software, 77, 1–17. 10.18637/jss.v077.i01

Xu, L., & Jordan, M. I. (1996). On Convergence Properties of the EM Algorithm for Gaussian Mixtures. Neural Computation, 8(1), 129–151. Neural Computation. 10.1162/neco.1996.8.1.129

Xu, Y., & Crouch, J. H. (2008). Marker-Assisted Selection in Plant Breeding: From Publications to Practice. Crop Science, 48(2), 391–407. 10.2135/cropsci2007.04.0191

Yang, J., Lee, S. H., Goddard, M. E., & Visscher, P. M. (2011). GCTA: A Tool for Genome-wide Complex Trait Analysis. American Journal of Human Genetics, 88(1), 76–82. 10.1016/j.ajhg.2010.11.011

Yao, Y., & Ochoa, A. (2023). Limitations of principal components in quantitative genetic association models for human studies. eLife, 12, e79238. 10.7554/eLife.79238

Yu, J., Pressoir, G., Briggs, W. H., Vroh Bi, I., Yamasaki, M., Doebley, J. F., McMullen, M. D., Gaut, B. S., Nielsen, D. M., Holland, J. B., Kresovich, S., & Buckler, E. S. (2006). A unified mixed-model method for association mapping that accounts for multiple levels of relatedness. Nature Genetics, 38(2), 203–208. 10.1038/ng1702

Zhang, J., Song, Q., Cregan, P. B., & Jiang, G.-L. (2016). Genome-wide association study, genomic prediction and marker-assisted selection for seed weight in soybean (Glycine max). TAG. Theoretical and Applied Genetics. Theoretische Und Angewandte Genetik, 129(1), 117–130. 10.1007/s00122-015-2614-x

Zhao, K., Tung, C.-W., Eizenga, G. C., Wright, M. H., Ali, M. L., Price, A. H., Norton, G. J., Islam, M. R., Reynolds, A., Mezey, J., McClung, A. M., Bustamante, C. D., & McCouch, S. R. (2011). Genome-wide association mapping reveals a rich genetic architecture of complex traits in Oryza sativa. Nature Communications, 2, 467. 10.1038/ncomms1467

Zhou, X., & Stephens, M. (2012). Genome-wide efficient mixed-model analysis for association studies. Nature Genetics, 44(7), 821–824. 10.1038/ng.2310

